# Targeted and untargeted nanopore sequencing approaches to profile the gut microbiota of mice infants exposed to ethanol *in utero*

**DOI:** 10.1101/2022.12.09.519725

**Authors:** Cristiano Pedroso-Roussado, Fergus Guppy, Nigel Brissett, Lucas Bowler, Joao Inacio

## Abstract

The gut microbiome plays a vital role in host homeostasis and understanding of its biology is essential for a better comprehension of the etiology of disorders such as foetal alcohol spectrum disorders. Here we assessed the effectiveness of targeted and untargeted (metagenomic) nanopore sequencing approaches to profile the gut microbiota of infant mice exposed to ethanol *in utero*. DNA extracts from the gut content of 12 infant mice exposed to ethanol in utero were analysed using one untargeted and two targeted (full-length 16S rRNA gene and the 16S-ITS-23S region of the ribosomal RNA operon) nanopore sequencing approaches. The targeting of the full-length 16S rRNA gene provided the most comprehensive analysis of the mouse gut microbiota. The differences in diversity between approaches were accounted by the sequencing target (p-value < 0.001). Faecalibaculum rodentium and Duncaniella sp. were the two most prevalent taxa detected using targeted sequencing approaches, while bacterial taxa were more evenly represented when using the metagenomic approach. Full-length 16S rRNA gene nanopore sequencing provides the most discriminatory microbiota compositional analysis of mice faecal samples. However, using nanopore sequencing approaches targeting the metagenome or different taxonomically-informative DNA region appears to introduce significant target-related biases.

**Importance:** Current nanopore approaches have not been standardized which may confound the biological interpretations of hight-throughput sequencing datasets. Additionally, nanopore sequencing still present a high error-rate compared to other more mature sequencing technologies, such as Illumina sequencing. These technological handicaps create the need to study and optimize nanopore sequencing approaches to answer biological questions, such as interrogations of the microbial composition and abundance of clinical and environmental samples. In this work, three nanopore sequencing approaches were designed and attempted to optimize fungal and bacterial profiling sequencing methodologies. Two targeted methods based on the bacterial 16S rRNA gene, and 16S-ITS-23S *rrn* operon region, and one untargeted shotgun/metagenomic approach were tested. Despite potential experimental and/or bioinformatical biases were found, the 16S rRNA gene-targeted nanopore sequencing was the most comprehensive approach to study the microbial composition of the infant mice gut microbiotas.

## Introduction

The study of the microbiomes has various experimental and data analytical challenges, due to the multidisciplinary nature of the area with input from microbiology, genomics, bioinformatics, epidemiology, among others. As a result, the consistency and commonality of procedures are not well established (1). Third-generation technologies such as nanopore sequencing whilst showing considerable promise still need developing to improve the reliability of sequencing outputs. To date two-way sequencing approaches have been employed to validate and/or complement nanopore sequencing studies together with more mature sequencing technologies, such as short-read Illumina sequencing. These experiments gain from the high-throughput resolution obtained by Illumina sequencing while increasing the sequencing coverage by the longer sequenced reads obtained by nanopore sequencing. Usually, nanopore sequencing achieves a higher number of taxa at the species level than Illumina sequencing (2), most likely explained by the capacity of nanopore sequencing reads to span multiple 16S rRNA variable regions (2).

Indeed, nanopore sequencing approaches rely on longer reads and reference databases are mostly composed of standard and shorter gene markers, such as fragments of the 16S rRNA gene or the ITS region (3, 4). Therefore, nanopore sequencing may have limited reliability in profiling unknown microbial communities, decreasing the confidence on the taxonomical composition obtained (5). Thus, targeting the V3-V4 hypervariable regions alone may overlook relevant genetic information necessary for further taxonomical separation at lower ranks, which could be solved by sequencing longer fragments encompassing multiple variable regions of 16S rRNA gene (2). Targeting the full-length 16S rRNA, the 16S-ITS-23S region of the *rrn* operon or performing a shotgun/metagenomic approach are potential ways to overcome these challenges (6, 7). However, the error-rate of nanopore sequencing is still too high to give total confidence in the approach, and further development both in sequencing chemistry and bioinformatic tools will undoubtedly resolve these issues by improving the accuracy of reads (8). Specifically, choosing the appropriate bioinformatic pipeline is crucial for a reliable output, which may represent a limitation in research settings where a better curated bioinformatic pipeline is necessary (8).

Here, two targeted and one untargeted nanopore sequencing approaches were optimized to profile the gut microbiota of 12 infant mice exposed to ethanol in utero. The sequencing was implemented using MinION system and the targeted approaches involved the sequencing of the full-length 16S rRNA gene and 16S-ITS-23S region of the *rrn* operon PCR-amplified amplicons. In the untargeted approach, a sequencing library was prepared directly from barcoded DNA templates extracted from the same infant mice samples.

## Results

In this study, total DNA was extracted from 12 infant mice faecal samples exposed to ethanol *in utero*. The aim was to deliberate the best bacterial genomic region for nanopore sequencing targeted approaches. Two targets were selected and compared – the full-length of the 16S rRNA gene (∼1,500 bp), and the whole *rrn* operon (16S rRNA gene-ITS-23S rRNA gene, with ∼4,500 bp). Additionally, the results obtained using these two targets were also compared with a shotgun/metagenomic microbial profiling approach (untargeted). The mean read quality of the nanopore sequencing runs was 10.1-10.8, and the total number of reads obtained per run was 374,000-1,454,000. One of the targeted approaches revealed an expected mean sequenced read length (1,500 bps for the full-length 16S rRNA gene). Interestingly, the average read length for the 16S-ITS-23S region of the operon was below the expected (3,708 bps *vs* 4,500 bps). Since the overall sequencing parameters were within the expected values (data not shown), it can be concluded that the shorter sequence read length observed was a direct consequence of a shorter DNA fragment. Despite the N50 of 6,353 bps obtained in the metagenomic approach, the average read length was 1,555 bps, which indicates that the DNA was fragmented throughout library preparation, leaving few longer fragments ready to be sequenced. The remaining nanopore sequencing quality parameters were in accordance with what was seen in the targeted approaches (data not shown).

### Influence of the nanopore sequencing approach in the microbial profiles

The three approaches were able to assign a different number of taxa showing the distinctive microbial profiling capacity of the genomic targets. At the phylum level, the full-length 16S rRNA gene-targeted nanopore sequencing approach assigned a total of 4 taxa, *Firmicutes* (74%), *Bacteroidetes* (22%), *Verrucomicrobia* (3%), and *Proteobacteria* (1%), while the 16S-ITS-23S region-targeted approach assigned 2, *Firmicutes* (51%) and *Bacteroidetes* (49%), and the metagenomic approach assigned 5, *Bacteroidetes* (5%), *Actinobacteria* (29%), *Proteobacteria* (11%), *Firmicutes* (5%), and *Verrucomicrobia* (1%). At the genus level, the full-length 16S rRNA gene-targeted approach assigned a total of 26 taxa, the most abundant being *Faecalibaculum* (28%), *Duncaniella* (11%), and *Muribaculum* (10%). The 16S-ITS-23S region-targeted approach assigned 16 taxa at the genus level, and the most abundant were *Faecalibaculum* (50%), *Duncaniella* (50%), *Muribaculum* (16%). The metagenomic approach assigned a total of 13 taxa at the genus level, and the most abundant were *Muribaculum* (29%), *Curtobacterium* (21%), and *Duncaniella* (18%). At the species level, the full-length 16S rRNA gene-targeted approach assigned a total of 31 taxa, while the 16S-ITS-23S region-targeted approach assigned 17, and the metagenomic approach assigned 13. The most abundant taxa at the species level were *Faecalibaculum rodentium* (28%), *Duncaniella muris* (11%), and *Muribaculum intestinale* (10%) in the 16S rRNA-based approach, *F. rodentium* (50%), *D. muris* (24%), and *M. intestinale* (16%) in the 16S-ITS-23S-based approach, and *M. intestinale* (29%), *Curtobacterium flaccumfaciens* (21%), and *D. muris* (18%) in the metagenomic approach (Figure 1).

**Figure 1.**
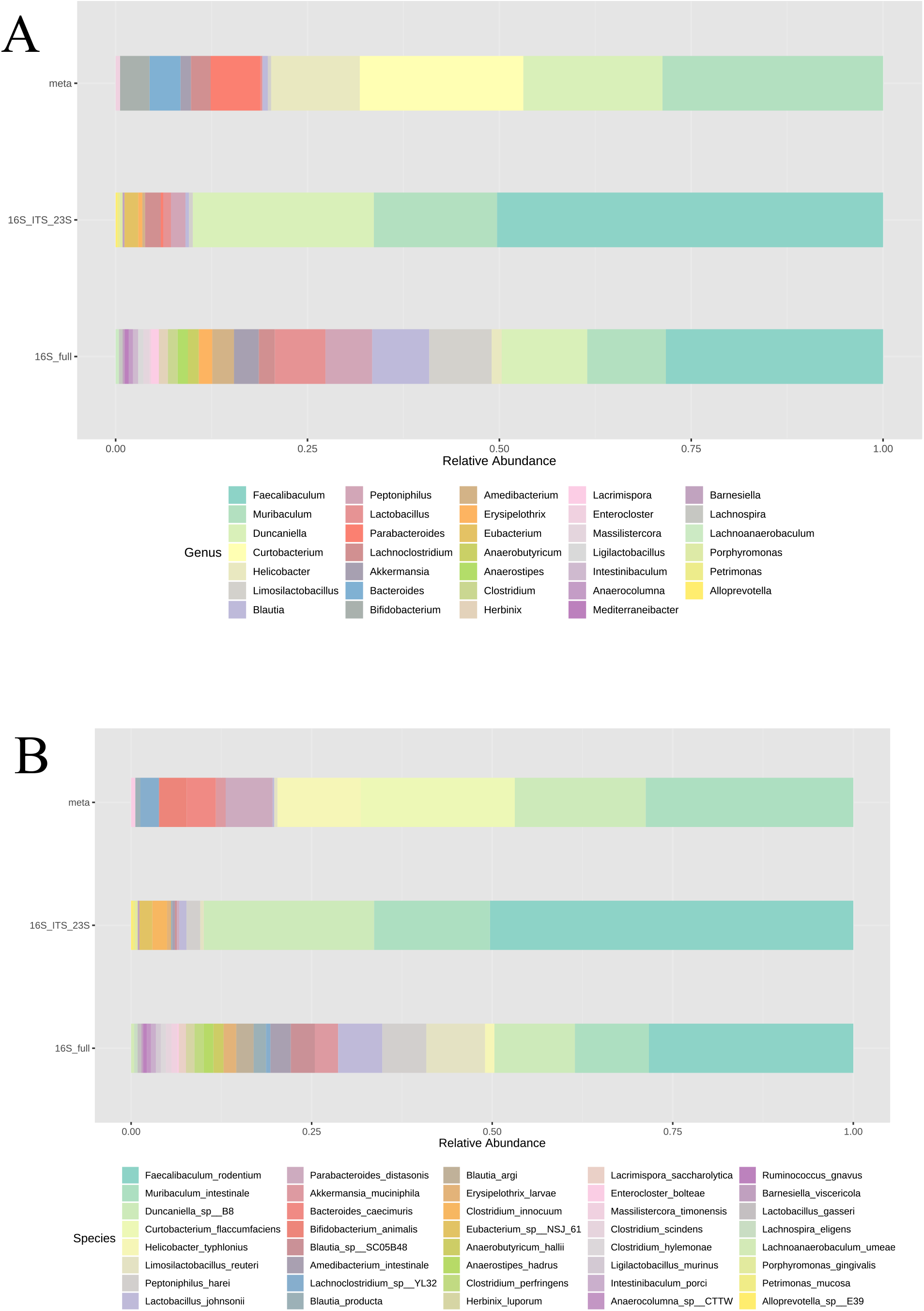
Bar chart showing the relative abundance of the combined taxa at the genus (A), and species level (C) detected by the 16S rRNA gene-, 16S-ITS-23S region-targeted, and untargeted shotgun/metagenomic nanopore sequencing approaches when 12 DNA templates extracted from infant mice faecal samples were pooled together.

The phylum *Verrucomicrobia* was only detected when the full-length 16S rRNA gene was used as genomic sequencing target. In the metagenomic approach, few reads were assigned to *Firmicutes*, and it was the only approach that detected *Proteobacteria*. Moreover, *C. flaccumfaciens* was only detected in the metagenomic approach with a relatively high abundance (21%).

### Assessment of potential biases and correlations found by targeted and untargeted approaches

Microbial profiles showed significant diversity differences within each infant mice faecal samples at the phylum, genus, and species level (Mann-Whitney U Test; p-value < 0.05). However, diversity similarities were also found within samples when comparing taxonomical genetic targets, specifically between full-length 16S rRNA gene and the 16S-ITS-23S operon region at the phylum level, and between the full-length 16S rRNA gene and the metagenomic approach at the genus and species level (Mann-Whitney U Test; p-value > 0.05, Figure 2).

**Figure 2.**
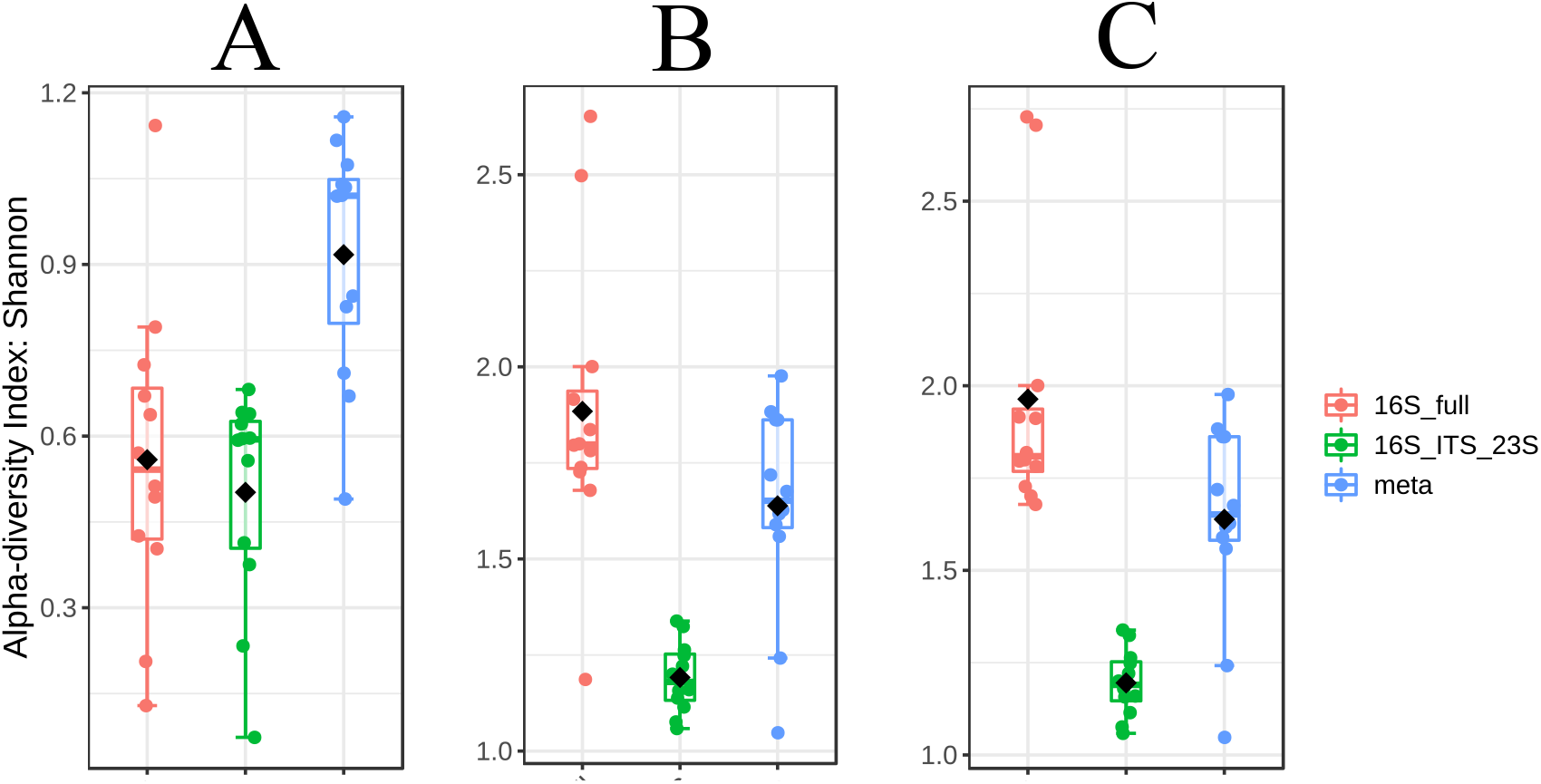
Boxplots displaying the combined alpha-diversity index at the phylum (A), genus (B), and species level (C), in the 16S rRNA gene-, 16S-ITS-23S region-targeted, and untargeted metagenomic/ shotgun nanopore sequencing approaches when 12 DNA templates extracted from infant mice faecal samples were pooled together.

Even though the microbial compositional differences observed between the 16S rRNA gene-targeted approach and the shotgun/metagenomic approach, the diversity found within their respective samples were not significantly different at the genus and species level. The untargeted and targeted approaches showed significant differences in the beta-diversity indices, representing the different microbial profiles obtained depending if the approach is targeted or untargeted. In the targeted approaches, nanopore sequencing was able to differentiate the diversity between samples at the genus and species level, however the level of dissimilarity was low (PERMANOVA R^2^ = 0.16-0.17; ANOSIM R = 0.25-0.27; p-value < 0.05). The dissimilarities in the beta-diversities were greater between the targeted and untargeted approaches (PERMANOVA R^2^ = 0.45-0.70; ANOSIM R = 0.69-0.81; p-value < 0.05). However, all groups showed significantly differences between themselves when analysed together (PERMANOVA R^2^ = 0.47-0.59; p-value < 0.05). Thus, these observations represent evidence of potential nanopore sequencing biases towards the experimental approach chosen.

To assess sequencing approach-based biases, correlations between assigned taxa and the 16S rRNA gene-, 16S-ITS-23S region-targeted, and untargeted shotgun/metagenomic nanopore sequencing approaches were analysed. *Limosillactobacillus* gen., *L. reuteri*, and *Blautia argi* were correlated with the full-length 16S rRNA gene marker, and phylum *Bacteroidetes*, genera *Duncaniella, Eubacterium, Muribaculum, Faecalibaculum, D. muris, Clostridium inocuum, Eubacterium* sp. NSJ 61, *M. intestinale*, and *F. rodentium*, correlated with the nanopore sequencing approach based on the 16S-ITS-23S operon region genomic target. Therefore, the potential nanopore sequencing biases may be related to the genomic target used for the amplicon generation and be significantly observed at phylum, genera, and species level alike.

When 16S rRNA gene-, 16S-ITS-23S region-targeted, and untargeted shotgun/metagenomic nanopore sequencing approaches were analysed together, six taxa were significantly correlated exclusively with one sequencing approach (Figure 3). *C. flaccumfaciens, Parabacteroides distasonis*, and *Bacteroides caecimuris* were significantly correlated with the untargeted shotgun/metagenomic nanopore sequencing approach (Figure 3A, 3B, 3C, and 3D), while *Peptinophilus harei* and *L. reuteri* were significantly correlated with the full-length 16S rRNA gene nanopore sequencing approach (Figure 3E and 3F).

**Figure 3.**
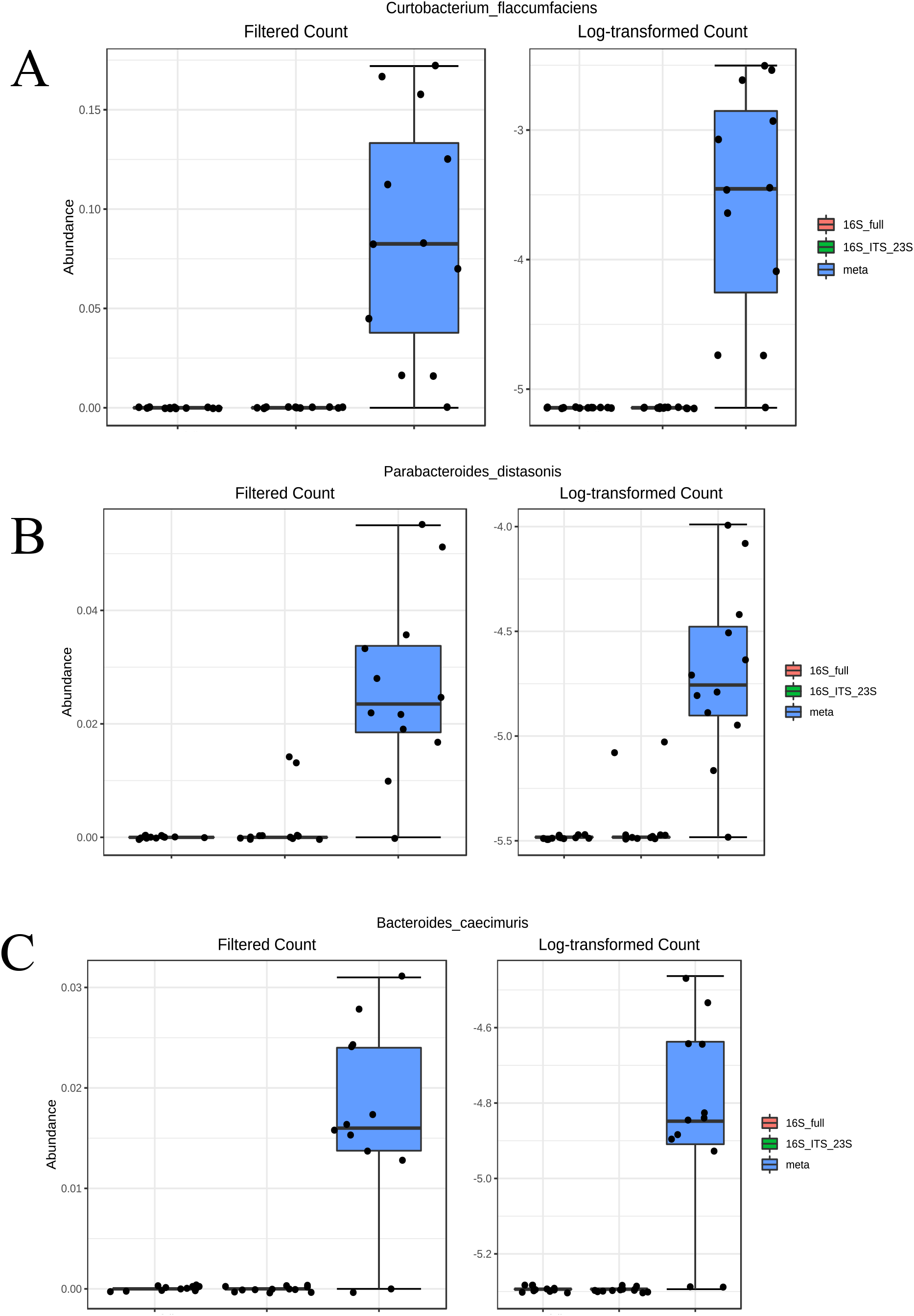
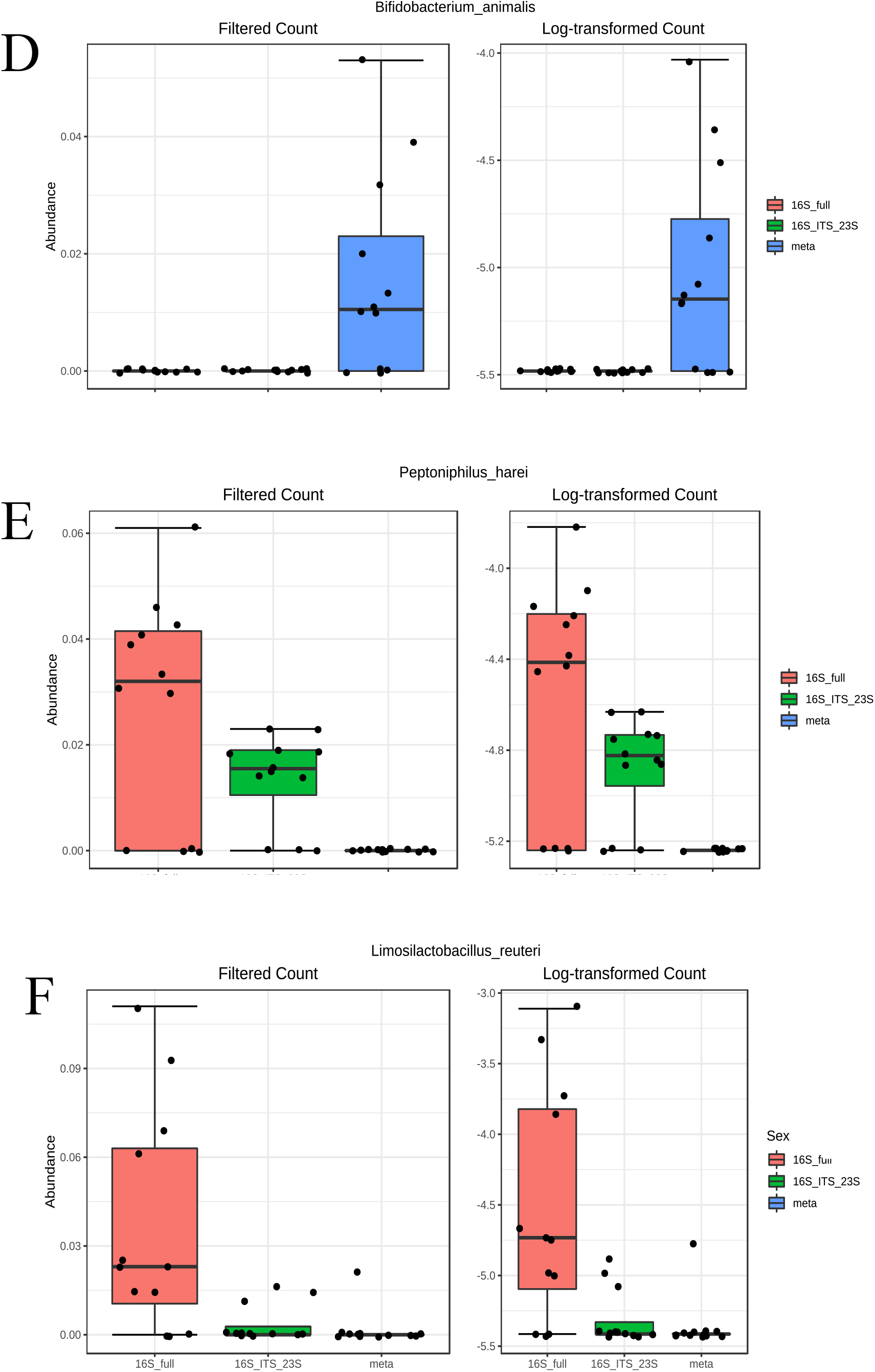
Boxplots displaying the relative abundances of taxa significantly correlated with one of the nanopore sequencing approaches tested. A, B, C, and D show the taxa correlated with the metagenomic/ shotgun nanopore sequencing approach; and E and F show the taxa correlated with the 16S rRNA gene-targeted nanopore sequencing approach. Left: raw relative abundances; Right: log-transformed relative abundances.

Therefore, the genomic target used for microbial profiling can induce biases at the compositional/population level, seen in the diversity indices, and at the individual level when nanopore sequencing approaches are attempted.

## Discussion

Despite the increased use of targeted sequencing approaches, one advantage of metagenomic approaches is the removal of upstream experimental steps, such as amplicon amplification by PCR, which may introduce amplification biases prior of library preparation (10) (11). In the metagenomic approach, most reads were not classified by WIMP (76.5%) and 35% of those classified were assigned to *Homo sapiens* (8.2% of total analysed reads) (data not shown). This observation might be explained by contamination throughout the sample handling and/or library preparations (12). However, due to the high basecalling error-rate of nanopore sequencing the hypothesis that reads assigned to *Homo sapiens* were misassigned, truly belonging to the mouse host, cannot be excluded; those reads were removed before further analyses were made. Besides the “DNA kitome” (13) (14), there are other type of contamination, such as reagents (15) (16), host material (17), and surrounding environment after sampling (18) (19). These contaminants would be sequenced alongside the desired sequencing library, potentially leading to biases in taxa abundances, sequencing coverage, specifically in library prepared with low biomass DNA (20). Additionally, the hypothesis of cross-contamination installed in the metagenomic reference databases cannot be fully discarded (17) (19) (20).

The low number of taxa detected by the 16S-ITS-23S region-targeted approach is most probably explained by the lower prevalence of sequences spanning the 16S-ITS-23S operon in the reference databases (21). The same observation can be made for the metagenomic approach, since the number and curation of whole-genomes present in the reference databases is not ideal (22). However, since longer reads were sequenced, there is a chance that some part of the read covered matched a more represented sequence in the reference database. Additionally, recent studies have been reporting a higher prevalence of taxa showing non-canonical (non-linked) 16S-ITS-23S DNA sequences (23) (24), and these non-linked arrangements have the potential to disturb taxonomical assignments when the 16S-ITS-23S region of the *rrn* operon is targeted for sequencing approaches (24). Still, some of these taxa were detected with a low prevalence and their detection may also be explained by basecalling errors occurred during nanopore sequencing.

The phylum *Verrucomicrobia* was only detected when the full-length 16S rRNA gene was used as genomic sequencing target. The only species assigned to this phylum was *Akkermansia muciniphila*. There is a lack of 16S-ITS-23S rRNA operon sequences representation in the databases and they may have dubious quality (Bowers *et al*., 2017, Shaiber and Eren, 2019). These observations and the possibility of occurring variants in the primer binding site (25) can explain why the targeted nanopore sequence approach based on the 16S-ITS-23S operon region was not able to assign *A. muciniphila* to any read. Moreover, the use of the 16S-ITS-23S operon region as a barcoding genetic marker is not always appropriate for microbial profiling (25). Since non-linked rRNA genes have been increasingly detected in environments that facilitates slower-growing or symbiotic taxa (23) (24), caution must be taken when choosing this type of barcodes for microbial profiling. Therefore, there is a relevant number of taxa that will be neglected if present in the samples under study, such as *Helicobacter, Rickettsia, Wolbachia, etc* (25). Up until now, unliked rRNA genes have been mostly reported in environments populated with slow-growing taxa like soils and sediments (28) (29). However, they might be more frequent and present in free-living taxa as well, including members of *Verrucomicrobia* (24). Interestingly, these taxa do not show slow growth patterns, low rRNA copy numbers, mutations’ fixation by genetic drift, small effective population sizes, nor reduced genome sizes (24). Indeed, these taxa are commonly abundant and ubiquitous in their environments and having non-linked rRNA genes could eventually confer a selective advantage by facilitating the production of heterogenous ribosomes with a diverse array of features (24). This speculative hypothesis is supported by recent reports that observed specialization in some rRNA loci detected in these taxa, which were translating genes involved in adaptation phenotypes related to temperature and nutrient shifts (30). Generally, it can be speculated that the different microbial profiles given by the two targeted nanopore approaches may be explained at least in part by the presence of non-linked 16S-ITS-23S DNA sequences as some taxa have separated 16S and 23S rRNA genes across the genome (24). Since there is an imbalance in the public databases that favours fast-growing and easily cultivated organisms (31) (32), it is expected that targeted sequencing approaches could find biases towards those taxa, neglecting others in possess of non-linked 16S-23S rRNA arrangements. However, this speculative explanation needs to be further studied in other organisms, such as in mice, since most, if not all, bacterial rRNA genes found in the human gut are linked (24).

Due to the underrepresentation of whole genomes in the databases, and the decreased chance of matching reads with the most represented genomic regions in the reference databases, it was expected that the metagenomic approach was unable to discriminate reads to lower taxonomical ranks than phylum level (22). Indeed, at the phylum level, the alpha-diversity index in the metagenomic approach was higher than the 16S rRNA gene-targeted approach, mostly due to the *Actinobacteria*, only detected in the metagenomic approach, and the higher abundance of *Proteobacteria* in comparison with the targeted approaches. At the genus and species level, the metagenomic approach revealed higher alpha-diversity indices when to compared to the 16S-ITS-23S region-targeted approach. This observation reveals that the metagenomic approach was more discriminatory and it was able to assign more taxa to the genus level than the mentioned targeted approach. This observation may be explained by the underrepresentation of 16S-ITS-23S operon regions sequences in the databases (26) (27). Regarding the diversity found within the samples analysed by the 16S-ITS-23S region-targeted approach, the low prevalence of correspondent sequences in the reference databases may undervalue the true diversity present in the infant mice gut. Indeed, targeting the 16S-ITS-23S operon region did not seem to improve the species level identification in comparison to the nanopore sequencing approach targeting the full-length 16S rRNA gene. This observation is supported by the least number of genera and species identified and, therefore, the smaller alpha- and beta-diversity indices found between the two targeted approaches. Thus, it is unlikely that the 16S-ITS-23S operon region can increase the taxonomic resolution, allowing strain level identification (6). Specifically, the distance between non-linked 16S and 23S rRNA genes in the genomes present in the datasets can span to 41 kilobases pairs or more, which constitutes a handicap for PCR amplification (24). Other attempts to microbial profile environmental samples with longer taxonomical barcodes have been successfully tried with other sequencing platforms, such as PacBio (21) (33) and Illumina synthetic long read sequencing, although limited to ∼2000 base pairs, but also with ONT (34) (35). Although the 16S-ITS-23S region-targeted nanopore sequencing approach here tested included the ITS region, the microbial profile obtained did not show increased species- nor strain-level composition and diversity, in accordance with the report by Karst and colleagues (34) (35), but contrary to others (36). Additionally, since some taxa (such *Lactobacillus reuteri* and *B. animalis* subsp. *animalis*) show dissimilarities within ITS copies of their strains and between each ITS copies in the same strain, the hypothesis of a better discriminatory power using this genomic barcode for targeted sequencing approaches is tempting but unlikely (21).

*Limosillactobacillus* gen., and *L. reuteri*, and *Blautia argi* were correlated with the full-length 16S rRNA gene marker, and *Bacteroidetes* phylum, genera *Duncaniella, Eubacterium, Muribaculum, Faecalibaculum, D. muris*, and *Clostridium inocuum, Eubacterium* sp. NSJ 61, *M. intestinale*, and *Faecalibaculum rodentium*, correlated with the nanopore sequencing approach based on the 16S-ITS-23S operon region genomic target. Therefore, the potential nanopore sequencing biases may be related to the genomic target used for the amplicon generation and be significantly observed at phylum, genera, and species level alike. Since nanopore sequencing is being increasingly explored for microbial profiling complex ecosystems, such as clinical and environmental samples, caution should be taken when targeted approaches are performed. The genomic target chosen may impose biases in the microbial composition consequently confounding the conclusions obtained by the experiments (36).

*C. flaccumfaciens, Parabacteroides distasonis*, and *B. caecimuris* were significantly correlated with the untargeted shotgun/metagenomic nanopore sequencing approach, while *Peptinophilus harei* and *Limosilactobacillus reuteri* were significantly correlated with the full-length 16S rRNA gene nanopore sequencing approach. Therefore, the genomic target used for microbial profiling can induce biases at the compositional/population level, seen in the diversity indices, and at the individual level when nanopore sequencing approaches are attempted. Consequently, the correlations found between these taxa and specific genomic targets lead to the conclusion that *C. flaccumfaciens, P. distasonis*, and *B. caecimuris*, can be considered “bias markers” for shotgun/metagenomic nanopore sequencing approaches, and *P. harei* and *L. reuteri* for the 16S rRNA gene-targeted nanopore sequencing approaches. Nevertheless, further investigation must be performed to elucidate the true meaning of the presence of these taxa and in which cases they may represent taxa erroneously assigned.

In conclusion, different gut microbiota profiles were obtained from the same samples when using different nanopore sequencing approaches. Both targeted approaches, based on the 16S rRNA gene and the 16S-ITS-23 region of the *rrn* operon, and untargeted approaches, yielded preferences towards specific taxa thus evidencing sequencing biases. In particular, *C. flaccumfaciens, P. distasonis, B. caecimuris*, and *B. animalis*, were correlated with the untargeted shotgun/metagenomic approach, and *P. harei*, and *L. reuteri* were correlated with the 16S rRNA gene-targeted approach. These results show that either upstream experimental steps, downstream bioinformatics analyses, or both, can introduce biases that are confounding the true microbial composition and abundances obtained from infant mice gut samples.

## Materials and Methods

### Ethical Statement

The mice experimental design was licenced under the Animals (Scientific Procedures) Act, 1986), with license number PPL P5686A14D. All experimental work was performed in accordance with the ARRIVE guidelines published by the National Centre for the Replacement Refinement & Reduction of Animals in Research. Animal Welfare and Review Board ethically approved all protocols and procedures.

### Animal husbandry

Three male and three female C57BL/6J mice (6 and 8 weeks old, respectively) were randomly paired into three groups. Each male randomly paired with one female and placed into an Optimice® cage to create three monogamous breeding pairs, with the female mice exposed to ethanol during pregnancy as part of other work investigating the effect of foetal alcohol exposure. A first batch of 12 infant mice exposed to ethanol *in utero* were used for the nanopore sequencing approaches optimization experiments. Cage and holding room environment were maintained constantly with controlled light intensity, relative humidity, room and cage temperature. To promote crepuscular behaviour, a 24 h lighting cycle of 12 h light followed by 12 h dark photoperiod was implemented in order to regulate both circadian rhythm and reproductive cycles. Enviro-Dry® fresh nesting was aided and preserved by microenvironment temperature. RM No.3 Special Diet Services (SDS) (5 g/day/mouse) was supplied to each group.

### Drug formulation and administration

Formulation of 5% ethanol solutions (1000 mL) were formed prior to drug administration and stored at 4 °C. To mimic average human ethanol consumption, for each group, 5% ethanol solution was orally administered from colony set-up onwards, by means of Optimice® internal bottle assembly and allowed fresh daily consumption. Offspring pups were weaned 23-28 days postnatal, separated and placed into gender specific boxes categorised with date of birth, wean date, and box origin.

### Infant mice gut DNA extraction and amplicons generation

DNA extraction was performed using around 300 mg of biological faecal material with QIAmp PowerFecal Pro DNA Kit, according to the manufacturer’s protocol (Qiagen, UK). The final supernatant containing the extracted DNA was used immediately or stored at -20 °C until further use. DNA quantity and quality (A_260/280_ and A_260/230_) were measured using a Qubit 4 Fluorometer (Thermofisher, USA), and with DeNovix DS-11 Spectrophotometer (DeNovix, USA), respectively. PCR was performed to amplify two targets, the full-length of the 16S rRNA gene (∼1,500 bp), and the whole *rrn* operon (16S rRNA gene-ITS-23S rRNA gene, with ∼4,500 bp). The primers pairs used were the 16S-27F – AGAGTTTGATCCTGGCTCAG, and the 16S-1492R – GGTTACCTTGTTACGACTT for the 16 rRNA gene. For the 16S-ITS-23S operon region., the same abovementioned 16S-27F was used together with the 23S-2241R – ACCRCCCCAGTHAAACT (6). The PCR mixture for the full-length 16S rRNA gene (25 μL total volume) contained between 0.2 and 1.6 ng of DNA template (or 3 μL of unquantified initial DNA extract), 1× Standard Taq Reaction Buffer, 0.5 mM of dNTPs, 0.4 μM of 16S-27F and 0.8 μM 16S-1492R, and 0.625 U of Taq DNA Polymerase (New England Biolabs, UK). The PCR thermal profile consisted of an initial denaturation of 30 s at 95 °C, followed by 30 cycles of 20 s at 95 °C, 20 s at 51 °C, 2 min at 68 °C, and a final step of 5 min at 72 °C. The PCR mixture for the full-length 16S-ITS-23 region of the *rrn* operon (25 μL total volume) contained between 0.2 and 1.6 ng of DNA template (or 3 μL of unquantified initial DNA), 1× Standard Taq Reaction Buffer, 0.5 mM of dNTPs, 1 μM of 16S-27F and 23S-2241R, and 0.625 U of Taq DNA Polymerase. The PCR thermal profile consisted of an initial denaturation of 30 s at 95 °C, followed by 30 cycles of 20 s at 95 °C, 20 s at 51 °C, 4 min and 30 s at 68 °C, and a final step of 5 min at 72 °C. The amplicons were purified with the AMPure XP beads (Beckman Coulter, UK) using a 0.5× and 0.40× ratio for the full-length 16S rRNA gene and the 16S-ITS-23S region of the *rrn* operon, respectively, according to manufacturer’s protocol. The quantity and quality of the purified DNA was measured by Qubit 4 Fluorometer, and with DeNovix DS-11 Spectrophotometer, respectively. If the PCR purified amplicons did not show A_260/280_ between 1.8-2.0 and A_260/230_ between 2.0-2.2, they were discarded, and the PCR reaction repeated.

### Libraries of 16S rRNA gene and 16S-ITS-23S operon amplicons

For the targeted approaches targeting the 16S rRNA gene and the 16S-ITS-23S region of the *rrn* operon, the Ligation Sequencing Kit (SQK-LSK 109, ONT, UK) was used in combination with PCR-Barcoding Expansion Kit (EXP-PBC001, ONT, UK) to tag each of the previously generated sample specific PCR amplicons and prepare the sequencing libraries, following the manufacturer’s protocol. The forward barcoding primer tag used was F – TTTCTGTTGGTGCTGATATTGC, and the reverse was R – ACTTGCCTGTCGCTCTATCTTC. Amplicons were quantified using the Qubit 4 Fluorometer and the volume was adjusted to begin the barcoding PCR with 100-200 fmol (0.5 nM) of the first PCR products, according to ONT instructions. PCR mixture for barcoding PCR (100 μL total volume) contained 100-200 fmol of each PCR product to be barcoded in 48 μL, 1× LongAmp® Taq 2× Master Mix (New England Biolabs, UK), and 2 μL of a unique barcode. The PCR thermal profile consisted of an initial denaturation of 3 min at 95 °C, followed by 13 cycles (for the 16S rRNA gene) or 15 cycles (for the 16S-ITS-23S region of the *rrn* operon) of 15 s at 95 °C, 15 s at 62 °C, 1 min and 30 s at 65 °C (for the 16S rRNA gene) or 4 min and 30 s at 65 °C (for the 16S-ITS-23S region of the *rrn* operon), and a final step of 1 min 30 s at 65 °C (for the 16S rRNA gene) or 4 min and 30 s (for the 16S-ITS-23S region of the *rrn* operon) at 65 °C. The barcoded amplicons were purified with the AMPure XP beads using a 0.5× and 0.40× ratio for the full-length 16S rRNA gene and the 16S-ITS-23S region of the *rrn* operon, respectively, according to the manufacturer’s protocol. Every tagged DNA sample were combined in a pool to reach 1 μg in a volume of 47 μL, 1 μL of DNA CS, 3.5 μL of NEBNext FFPE DNA Repair Buffer (New England Biolabs, UK), 2 μL of NEBNext FFPE DNA Repair Mix (New England Biolabs, UK), 3.5 μL Ultra II End-prep reaction buffer, and 3 μL of Ultra-prep enzyme mix (New England Biolabs, UK), and completed with molecular biology grade water. The mixture was incubated in a thermal cycler at 20 °C for 5 min and 65 °C for 5 min. A purification step using 1× Agencourt AMPure XP beads was performed. Briefly, the mixture was incubated at room temperature with gentle mixing for 5 min, washed twice with 200 μL fresh 70% ethanol, pellet allowed to dry at 37 °C until a light brown colour was observed before DNA being eluted in 61 μL nuclease-free water, and incubated at room temperature for 30 min or more. A 1 μL aliquot was quantified by fluorometry using Qubit 4 Fluorometer. For the adapter ligation step, the total 60 μL of end-prepped DNA from the previous step were added in a mix containing 25 μL of Ligation Buffer LNB (NEBNext Quick Ligation Module, New England Biolabs, UK), 10 μL NEBNext Quick T4 DNA Ligase (New England Biolabs, UK), and 5 μL of Adapter Mix (ONT, UK), and incubated at room temperature for 10 min. A purification step using 0.4× Agencourt AMPure XP beads was performed. Briefly, the mixture was incubated at room temperature with gentle mixing for 5 min, washed twice with 250 μL of Short Fragment Buffer (ONT, UK pellet allowed to dry at 37 °C until a light brown colour was observed before DNA being resuspended in 15 μL of Elution Buffer (ONT, UK), and incubated at room temperature for 30 min or more. The mixture was retained. A 1 μL aliquot was quantified by fluorometry using Qubit 4 Fluorometer and quality measured with DeNovix DS-11 Spectrophotometer to ensure that no contaminants were present before the library loading step. If the DNA sequencing library did not show A_260/280_ between 1.8-2.0 and A_260/230_ between 2.0-2.2, it was discarded, and the protocol repeated.

### Library for the shotgun/metagenomic nanopore sequencing approach

For the untargeted shotgun/metagenomic nanopore sequencing approach the same sequencing kit was used in combination with Native Barcoding Kit (EXP-NBD104, ONT, UK) to tag each DNA template extracted from the gut samples with an unique barcode, following the manufacturer’s protocol. The forward barcoding primer tag used was F – TTTCTGTTGGTGCTGATATTGC, and the reverse was R – ACTTGCCTGTCGCTCTATCTTC. Every tagged DNA sample were pooled together to reach 1 μg in a volume of 48 μL containing 3.5 μL of NEBNext FFPE DNA Repair Buffer, 2 μL of NEBNext FFPE DNA Repair Mix, 3.5 μL Ultra II End-prep reaction buffer, and 3 μL of Ultra-prep enzyme mix, and completed with molecular biology grade water. The mixture was incubated in a thermal cycler at 20 °C for 5 min and 65 °C for 5 min. A purification step using 1× Agencourt AMPure XP beads was performed. Briefly, the mixture was incubated at room temperature with gentle mixing for 5 min, washed twice with 200 μL fresh 70% ethanol, pellet allowed to dry at 37 °C until a light brown colour was observed before DNA being resuspended in 25 μL nuclease-free water and incubated at room temperature for 1 h or more. A 1 μL aliquot was quantified by fluorometry using Qubit 4 Fluoremeter to ensure that at least 700 ng of DNA was retained. For barcode ligation, 500 ng of end-prepped DNA was added separately to 2.5 μL of a unique Native Barcode, and 25 μL of NEB Blunt/TA Ligase Master Mix (New England Biolabs, UK). The mixture was incubated at room temperature for 10 min. A purification step using 1× Agencourt AMPure XP beads was performed. Briefly, the mixture was incubated at room temperature with gentle mixing for 5 min, washed twice with 200 μL fresh 70% ethanol, pellet allowed to dry at 37 °C until a light brown colour was observed before DNA being resuspended in 26 μL nuclease-free water, and incubated at room temperature for 1 h or more. A 1 μL aliquot was quantified by fluorometry using Qubit 4 Fluorometer. Equimolar concentrations of each native barcoded sample were pooled together assuring that a total 700 ng of DNA was guaranteed. A 1 μL aliquot was quantified by fluorometry using Qubit 4 Fluorometer. For the adapter ligation step, 65 μL containing 700 ng of pooled end-prepped native barcoded DNA from the previous step were added in a mix containing 5 μL of Adapter Mix (AMII, ONT, UK), 20 μL NEBNext 5× Quick Ligation Reaction Buffer (New England Biolabs, UK), and 10 μL of Quick T4 DNA Ligase, and incubated at room temperature for 10 min. A purification step using 0.5× Agencourt AMPure XP beads was performed. Briefly, the mixture was incubated at room temperature with gentle mixing for 5 min, washed twice with 250 μL of Short Fragment Buffer, pellet allowed to dry at 37 °C until a light brown colour was observed before DNA being eluted in 15 μL of Elution Buffer, and incubated at room temperature for 30 min or more. The mixture was retained. A 1 μL aliquot was quantified by fluorometry using Qubit 4 Fluorometer and quality measured with DeNovix DS-11 Spectrophotometer to ensure no contaminants were present before the library loading step. If the DNA sequencing library did not show A_260/280_ between 1.8-2.0 and A_260/230_ between 2.0-2.2, it was discarded, and the protocol repeated.

### MinION sequencing, basecalling. and bioinformatics analysis

Nanopore sequencing was performed using MinION MK1B sequencer (ONT, UK) and flowcells R9.4.1 (FLOW-MIN106D, ONT, UK) using MinKNOW™ software v.21.02.1 (ONT, UK). After the MinION Flow Cell was setup accordingly to manufacturer’s instructions, the pre-sequencing mix was prepared accordingly to manufacturer’s protocol. Once the DNA sequencing library was loaded, the 24 h sequencing run was initiated. For the targeted sequencing approach using the full-length 16S rRNA gene, two MinION Flow Cells were used so that 6 barcoded DNA samples were sequenced per sequencing run. In the targeted sequencing approach using the 16S-ITS-23S operon region and the shotgun/metagenomic nanopore sequencing approach, a single flowcell was used with 12 barcoded DNA samples. All nanopore sequencing experiments ran for 24 consecutive hours. Basecalling was performed with *guppy* standalone tool (https://nanoporetech.com) from the command line using the *fast basecalling* mode, with the quality score filter of 8, and adapters’ trimming mode *on*. Quality controls were performed using *nanoplot* tool (38) with the length filters parameters of maximum length at 2000 and minimum at 1000 for the 16S rRNA gene-targeted approach, maximum length at 5000 and minimum at 1000 for the 16S-ITS-23S region of the *rrn* operon-targeted approach, and minimum at 300, with no maximum length filter for the shotgun/metagenomic approach. WIMP (version 2021.11.26) was used for the classification assignments steps.

### Statistical analysis

For comparative analyses, non-normalized relative frequencies were used as input for MicrobiomeAnalyst software (39), and only taxa with relative abundances above 1% were considered. Appropriate statistical tests are described in the text.

## Acknowledgements

This research was supported by the DTA3/COFUND PhD Fellowship Programme (funding from the European Union’s Horizon 2020 research and innovation programme under the Marie Skłodowska-Curie grant agreement No. 801604). The funder had no role in study design, data collection and interpretation, or the decision to submit the work for publication.

Conceptualization, C.P-R., F.G., L.B., and J.I.; Methodology C.P-R, F.G., L.B., and J.I.;

Investigation, C.P-R.; Mice experimental work, N.B.; Supervision, F.G., L.B., and J.I.;

Writing – original draft, C.P-R.; Writing – review & editing, F.G., N.B., L.B., and J.I.

## Appendixes

## References

1. Simoneau J, Dumontier S, Gosselin R, Scott MS. 2021. Current RNA-seq methodology reporting limits reproducibility. Brief Bioinform. 22(1):140–145.

2. Shin J, Lee S, Go MJ, Lee SY, Kim SC, Lee CH, Cho BK. 2016. Analysis of the mouse gut microbiome using full-length 16S rRNA amplicon sequencing. Sci Rep. 6:29681.

3. Markey L, Hooper A, Melon LC, Baglot S, Hill MN, Maguire J, Kumamoto CA. 2020. Colonization with the commensal fungus Candida albicans perturbs the gut-brain axis through dysregulation of endocannabinoid signaling. Psychoneuroendocrinology. 121:104808.

4. Matsuo Y, Komiya S, Yasumizu Y, Yasuoka Y, Mizushima K, Takagi T, Kryukov K, Fukuda A, Morimoto Y, Naito Y, Okada H, Bono H, Nakagawa S, Hirota K. 2021. Full-length 16S rRNA gene amplicon analysis of human gut microbiota using MinION™ nanopore sequencing confers species-level resolution. BMC Microbiol. 21(1):35.

5. Santos A, van Aerle R, Barrientos L, Martinez-Urtaza J. 2020. Computational methods for 16S metabarcoding studies using Nanopore sequencing data. Comput Struct Biotechnol J.18:296–305.

6. Cuscó A, Catozzi C, Viñes J, Sanchez A, Francino O. 2018. Microbiota profiling with long amplicons using Nanopore sequencing: full-length 16S rRNA gene and the 16S-ITS-23S of the rrn operon. F1000Res. 7:1755.

7. Jeon S, Kim H, Kim J, Seol D, Jo J, Choi Y, Cho S, Kim H. 2022. Positive Effect of Lactobacillus acidophilus EG004 on Cognitive Ability of Healthy Mice by Fecal Microbiome Analysis Using Full-Length 16S-23S rRNA Metagenome Sequencing. Microbiol Spectr. 10(1):e0181521.

8. Leggett RM, Alcon-Giner C, Heavens D, Caim S, Brook TC, Kujawska M, Martin S, Peel N, Acford-Palmer H, Hoyles L, Clarke P, Hall LJ, Clark MD. 2020. Rapid MinION profiling of preterm microbiota and antimicrobial-resistant pathogens. Nat Microbiol. 5(3):430–442.

9. Shafin K, Pesout T, Lorig-Roach R, Haukness M, Olsen HE, Bosworth C, Armstrong J, Tigyi K, Maurer N, Koren S, Sedlazeck FJ, Marschall T, Mayes S, Costa V, Zook JM, Liu KJ, Kilburn D, Sorensen M, Munson KM, Vollger MR, Monlong J, Garrison E, Eichler EE, Salama S, Haussler D, Green RE, Akeson M, Phillippy A, Miga KH, Carnevali P, Jain M, Paten B. 2020. Nanopore sequencing and the Shasta toolkit enable efficient de novo assembly of eleven human genomes. Nat Biotechnol.38(9):1044–1053.

10. Nilsson RH, Anslan S, Bahram M, Wurzbacher C, Baldrian P, Tedersoo L. 2019. Mycobiome diversity: high-throughput sequencing and identification of fungi. Nat Rev Microbiol. 17(2):95–109.

11. Nearing JT, Comeau AM, Langille MGI. 2021. Identifying biases and their potential solutions in human microbiome studies. Microbiome. 9(1):113.

12. Eisenhofer R, Minich JJ, Marotz C, Cooper A, Knight R, Weyrich LS. 2019. Contamination in Low Microbial Biomass Microbiome Studies: Issues and Recommendations. Trends Microbiol. 27(2):105–117.

13. Salter SJ, Cox MJ, Turek EM, Calus ST, Cookson WO, Moffatt MF, Turner P, Parkhill J, Loman NJ, Walker AW. 2014. Reagent and laboratory contamination can critically impact sequence-based microbiome analyses. BMC Biol. 12:87.

14. Olomu IN, Pena-Cortes LC, Long RA, Vyas A, Krichevskiy O, Luellwitz R, Singh P, Mulks MH. 2020. Elimination of “kitome” and “splashome” contamination results in lack of detection of a unique placental microbiome. BMC Microbiol. 20(1):157.

15. Kulakov LA, McAlister MB, Ogden KL, Larkin MJ, O’Hanlon JF. 2002. Analysis of bacteria contaminating ultrapure water in industrial systems. Appl Environ Microbiol. 68(4):1548–55.

16. Olm MR, Butterfield CN, Copeland A, Boles TC, Thomas BC, Banfield JF. 2017. The Source and Evolutionary History of a Microbial Contaminant Identified Through Soil Metagenomic Analysis. mBio. 8(1):e01969–16.

17. Ames SK, Gardner SN, Marti JM, Slezak TR, Gokhale MB, Allen JE. 2015. Using populations of human and microbial genomes for organism detection in metagenomes. Genome Res. 25(7):1056–67.

18. Lusk RW. 2014. Diverse and widespread contamination evident in the unmapped depths of high throughput sequencing data. PLoS One. 9(10):e110808.

19. Gruber K. 2015. Here, there, and everywhere: From PCRs to next-generation sequencing technologies and sequence databases, DNA contaminants creep in from the most unlikely places. EMBO Rep. 16(8):898–901.

20. Lu J, Salzberg SL. 2018. Removing contaminants from databases of draft genomes. PLoS Comput Biol. 14(6):e1006277.

21. Martijn J, Lind AE, Schön ME, Spiertz I, Juzokaite L, Bunikis I, Pettersson OV, Ettema TJG. 2019. Confident phylogenetic identification of uncultured prokaryotes through long read amplicon sequencing of the 16S-ITS-23S rRNA operon. Environ Microbiol. 21(7):2485–2498.

22. Breitwieser FP, Lu J, Salzberg SL. 2019. A review of methods and databases for metagenomic classification and assembly. Brief Bioinform. 20(4):1125–1136.

23. Ahn H, Seol D, Cho S, Kim H, Kwak W. 2020. Enhanced Symbiotic Characteristics in Bacterial Genomes with the Disruption of rRNA Operon. Biology (Basel). 9(12):440.

24. Brewer TE, Albertsen M, Edwards A, Kirkegaard RH, Rocha EPC, Fierer N. 2020. Unlinked rRNA genes are widespread among bacteria and archaea. ISME J. 14(2):597–608.

25. Seol D, Lim JS, Sung S, Lee YH, Jeong M, Cho S, Kwak W, Kim H. 2022. Microbial Identification Using rRNA Operon Region: Database and Tool for Metataxonomics with Long-Read Sequence. Microbiol Spectr. 10(2):e0201721.

26. Bowers RM, Kyrpides NC, Stepanauskas R, Harmon-Smith M, Doud D, Reddy TBK, Schulz F, Jarett J, Rivers AR, Eloe-Fadrosh EA, Tringe SG, Ivanova NN, Copeland A, Clum A, Becraft ED, Malmstrom RR, Birren B, Podar M, Bork P, Weinstock GM, Garrity GM, Dodsworth JA, Yooseph S, Sutton G, Glöckner FO, Gilbert JA, Nelson WC, Hallam SJ, Jungbluth SP, Ettema TJG, Tighe S, Konstantinidis KT, Liu WT, Baker BJ, Rattei T, Eisen JA, Hedlund B, McMahon KD, Fierer N, Knight R, Finn R, Cochrane G, Karsch-Mizrachi I, Tyson GW, Rinke C; Genome Standards Consortium, Lapidus A, Meyer F, Yilmaz P, Parks DH, Eren AM, Schriml L, Banfield JF, Hugenholtz P, Woyke T. 2017. Minimum information about a single amplified genome (MISAG) and a metagenome-assembled genome (MIMAG) of bacteria and archaea. Nat Biotechnol. 35(8):725–731. Erratum in: Nat Biotechnol. 36(2):196. Erratum in: Nat Biotechnol. 36(7):660.

27. Shaiber A, Eren AM. 2019. Composite Metagenome-Assembled Genomes Reduce the Quality of Public Genome Repositories. mBio. 10(3):e00725–19.

28. Vieira-Silva S, Rocha EP. 2010. The systemic imprint of growth and its uses in ecological (meta)genomics. PLoS Genet. 6(1):e1000808.

29. Brown CT, Olm MR, Thomas BC, Banfield JF. 2016. Measurement of bacterial replication rates in microbial communities. Nat Biotechnol. 34(12):1256–1263.

30. Song W, Joo M, Yeom JH, Shin E, Lee M, Choi HK, Hwang J, Kim YI, Seo R, Lee JE, Moore CJ, Kim YH, Eyun SI, Hahn Y, Bae J, Lee K. 2019. Divergent rRNAs as regulators of gene expression at the ribosome level. Nat Microbiol. 4(3):515–526.

31. Vartoukian SR, Palmer RM, Wade WG. 2010. Strategies for culture of ‘unculturable’ bacteria. FEMS Microbiol Lett. 309(1):1–7.

32. Zhi XY, Zhao W, Li WJ, Zhao GP. 2012. Prokaryotic systematics in the genomics era. Antonie Van Leeuwenhoek. 101(1):21–34.

33. Jamy M, Foster R, Barbera P, Czech L, Kozlov A, Stamatakis A, Bending G, Hilton S, Bass D, Burki F. 2020. Long-read metabarcoding of the eukaryotic rDNA operon to phylogenetically and taxonomically resolve environmental diversity. Mol Ecol Resour. 20(2):429–443.

34. Karst SM, Ziels RM, Kirkegaard RH, Sørensen EA, McDonald D, Zhu Q, Knight R, Albertsen M. 2021. High-accuracy long-read amplicon sequences using unique molecular identifiers with Nanopore or PacBio sequencing. Nat Methods.18(2):165–169.

35. Karst SM, Dueholm MS, McIlroy SJ, Kirkegaard RH, Nielsen PH, Albertsen M. 2018. Retrieval of a million high-quality, full-length microbial 16S and 18S rRNA gene sequences without primer bias. Nat Biotechnol. 36(2):190–195.

36. de Oliveira Martins L, Page AJ, Mather AE, Charles IG. 2020. Taxonomic resolution of the ribosomal RNA operon in bacteria: implications for its use with long-read sequencing. NAR Genom Bioinform. 2(1):qz016.

37. Delahaye C, Nicolas J. 2021. Sequencing DNA with nanopores: Troubles and biases. PLoS One.16(10):e0257521.

38. De Coster W, D’Hert S, Schultz DT, Cruts M, Van Broeckhoven C. 2018. NanoPack: visualizing and processing long-read sequencing data. Bioinformatics. 34(15):2666–2669.

39. Chong J, Liu P, Zhou G, Xia J. 2020. Using MicrobiomeAnalyst for comprehensive statistical, functional, and meta-analysis of microbiome data. Nat Protoc. 15(3):799–821.

